# A Weighted Two-stage Sequence Alignment Framework to Identify DNA Motifs from ChIP-exo Data

**DOI:** 10.1101/2023.04.06.535915

**Authors:** Yang Li, Yizhong Wang, Cankun Wang, Anne Fennell, Anjun Ma, Jing Jiang, Zhaoqian Liu, Qin Ma, Bingqiang Liu

## Abstract

Identifying precise transcription factor binding sites (TFBS) or regulatory DNA motifs plays a fundamental role in researching transcriptional regulatory mechanisms in cells and in helping construct regulatory networks. Current algorithms developed for motif searching focus on the analysis of ChIP-enriched peaks but are not able to integrate the ChIP signal in nucleotide resolution. We present a weighted two-stage alignment tool (TESA). Our framework implements an analysis workflow from experimental datasets to TFBS prediction results. It employs a binomial distribution model and graph searching model with ChIP-exonuclease (ChIP-exo) reads depth and sequence data. TESA can effectively measure the possibility for each position to be an actual TFBS in a given promoter sequence and predict statistically significant TFBS sequence segments. The algorithm substantially improves prediction accuracy and extends the scope of applicability of existing approaches. We apply the framework to a collection of 20 ChIP-exo datasets of E. coli from proChIPdb and evaluate the prediction performance through comparison with three existing programs. The performance evaluation against the compared programs indicates that TESA is more accurate for identifying regulatory motifs in prokaryotic genomes.

## INTRODUCTION

Sequence-binding proteins are a diverse group of proteins that bind to specific sequences of DNA or RNA, often playing a crucial role in regulating gene expression and other cellular processes, which include transcription factors (TFs), RNA binding proteins, and chromatin-associated proteins, etc. ^1-8^. TFs are proteins that exert a pivotal impact in gene expression by binding to specific DNA sequences and either promoting or inhibiting transcription^3^. Understanding TF binding can help identify key regulatory elements in the genome, such as enhancers and promoters, shedding light on the complex network of interactions between TFs and other proteins involved in transcriptional regulation^9-11^. Short DNA sequences that are recognized and bound by a specific TF can be characterized by TF motifs. These TF motifs can be used to develop computational models of gene regulation as well as to design experiments to test the role of specific TFs in gene expression^9,10^.

ChIP-seq (chromatin immunoprecipitation sequencing) is a method to identify the genomic locations where a particular protein, such as a TF or a histone modification, is bound to DNA^12^. By providing a genome-wide map of protein-DNA interactions, ChIP-seq has enabled the identification of novel TF binding sites (TFBSs) and the refinement of existing TF motif models. There are a great many motif finding tools that are applicable for ChIP-seq, including DREME^13^, MEME-ChIP^7^, Weeder^14^, ProSampler^1^, and STREME^13^, etc. These tools vary in their algorithms, input requirements, and outputs, and can be used for different types of motif discovery and analysis. The motif finding algorithm, MEME, is a popular tool for discovering overrepresented motifs in a set of DNA sequences^15^. MEME uses the expectation-maximization (EM) algorithm to estimate a probability distribution for the occurrence of each motif in the input sequences. The algorithm starts with an initial set of motifs, and then iteratively improves the motif models by refining the estimated probability distributions for each motif and identifying new candidate motifs based on the distribution of sequence patterns in the input sequences. DREME is designed to identify motifs that are statistically overrepresented in a set of DNA sequences, using a discriminative approach that distinguishes between positive and negative examples^13^. DREME is particularly useful for discovering motifs in large and complex datasets, where traditional motif discovery algorithms may be limited by the size and complexity of the input sequences. MEME-ChIP uses MEME and DREME in a complementary manner to improve the accuracy and specificity of motif discovery in ChIP-seq data. By combining these two approaches, MEME-ChIP can identify motifs that are both highly enriched and statistically significant, providing a more accurate and comprehensive view of the TF binding motifs in ChIP-seq data. STREME is a motif finding algorithm that is used in place of DREME in the current MEME-ChIP pipeline^13^. Like DREME, STREME uses regular expression patterns to represent motifs, but it employs a more efficient algorithm for motif discovery based on suffix trees, making it well-suited for analyzing large and complex datasets. Similar to STREME, Weeder uses a suffix tree data structure to efficiently identify frequent *k*-mers (short DNA sequences of length *k*) and extend them into longer motifs^14^. ProSampler is a discriminative motif finding algorithm that uses a probabilistic sampling approach to identify overrepresented motifs in a set of DNA sequences^1^. ProSampler identifies motif lengths without using exhaustive search within a range. ProSampler is designed to be efficient and scalable, making it well-suited for analyzing large and complex datasets.

ChIP-exo (ChIP with exonuclease treatment and high-throughput sequencing) is a higher-resolution version of ChIP-seq that uses an exonuclease enzyme to precisely cleave the DNA fragments bound to the protein of interest, resulting in a more accurate mapping of protein-DNA interactions^16,17^. ChIP-exo provides higher resolution of protein-DNA interactions than ChIP-seq but also presents three challenges due to its more complex data analysis. First, ChIP-exo data has higher sensitivity to experimental conditions than ChIP-seq. Second, unlike ChIP-seq, the binding sites in ChIP-exo may be located at the ends of the signal peaks, conflicting with the assumption of most algorithms on ChIP-seq that the binding sites of a TF are located at the center of the ChIP-seq peaks. Third, ChIP-exo can produce sharper peaks than ChIP-seq, which makes it more difficult to accurately distinguish between adjacent binding events. To cope with these challenges, we put forth a weighted two-stage alignment (TESA) tool on the basis of BoBro^18^. TESA could overcome the three challenges using the following strategies. First, TESA enhances the signal-to-noise ratio by clustering highly similar potential TFBSs via a graph model and optimizing motif lengths using a bookend model. Second, TESA scores sequence segments based on a binomial distribution which integrates the positional sequencing coverage instead of assuming that TFs are apt to binding the peak center. Third, TESA uses a binomial test to determine whether two clusters of potential TFBSs should be combined or treated as separate entities.

TESA is tested on 20 ChIP-exo datasets of E. *coli* from proChIPdb and can identify both known motifs and novel ones^19^. When compared to three state-of-the-art methods, BoBro, MEME-ChIP, and Weeder, TESA is found to have superior performance in distinguishing TFBSs and non-TFBSs, evaluated by four metrics. Additionally, the motifs discovered by TESA showcase significant similarity with the ones curated in the DPInteract database^20^, evaluated by TOMTOM^21^.

## MATERIALS AND METHODS

### Data acquisition

A total of 20 ChIP-exo datasets of E. *coli* are used in this study. For each dataset, we regard all its peaks as positive sequences with label ‘1’. We generate negative sequences with label ‘0’ by randomly shuffling all bases within a positive sequence^22^. The negative sequences do not contain TFBSs but possess the same GC contents as the positive ones. The union set of positive and negative sequences constitutes the sequences for benchmarking analysis.

### Overview of the TESA pipeline

The basic idea of TESA can be explained as follows (**Figure 1**). After data pre-processing via MACE and BEDTOOLs^23,24^, TESA detects potential TFBSs by a scoring approach using binomial distribution for sequence segments of a fixed length in an exhaustive manner on each sequence. Then TESA constructs a graph, in which vertices represent sequence segments and an unweighted edge connecting two vertices indicates a highly ranked similarity between them among all pairs of sequence segments between two sequences. TESA identifies dense subgraphs as the seed for graph clustering. Then, TESA performs graph clustering based on seeds, leading to vertex clusters, each of which corresponds to a preliminary motif. Afterwards, by assembling clusters with a significant overlap evaluated by binomial distribution, TESA optimizes the lengths of preliminary motifs using a bookend model. We call the sequence segments corresponding to the assembled clusters as motif seeds. As the last step, TESA refines the sequence segments for each motif, by scoring them using the motif profile built from the motif seeds.

**Figure 1.**
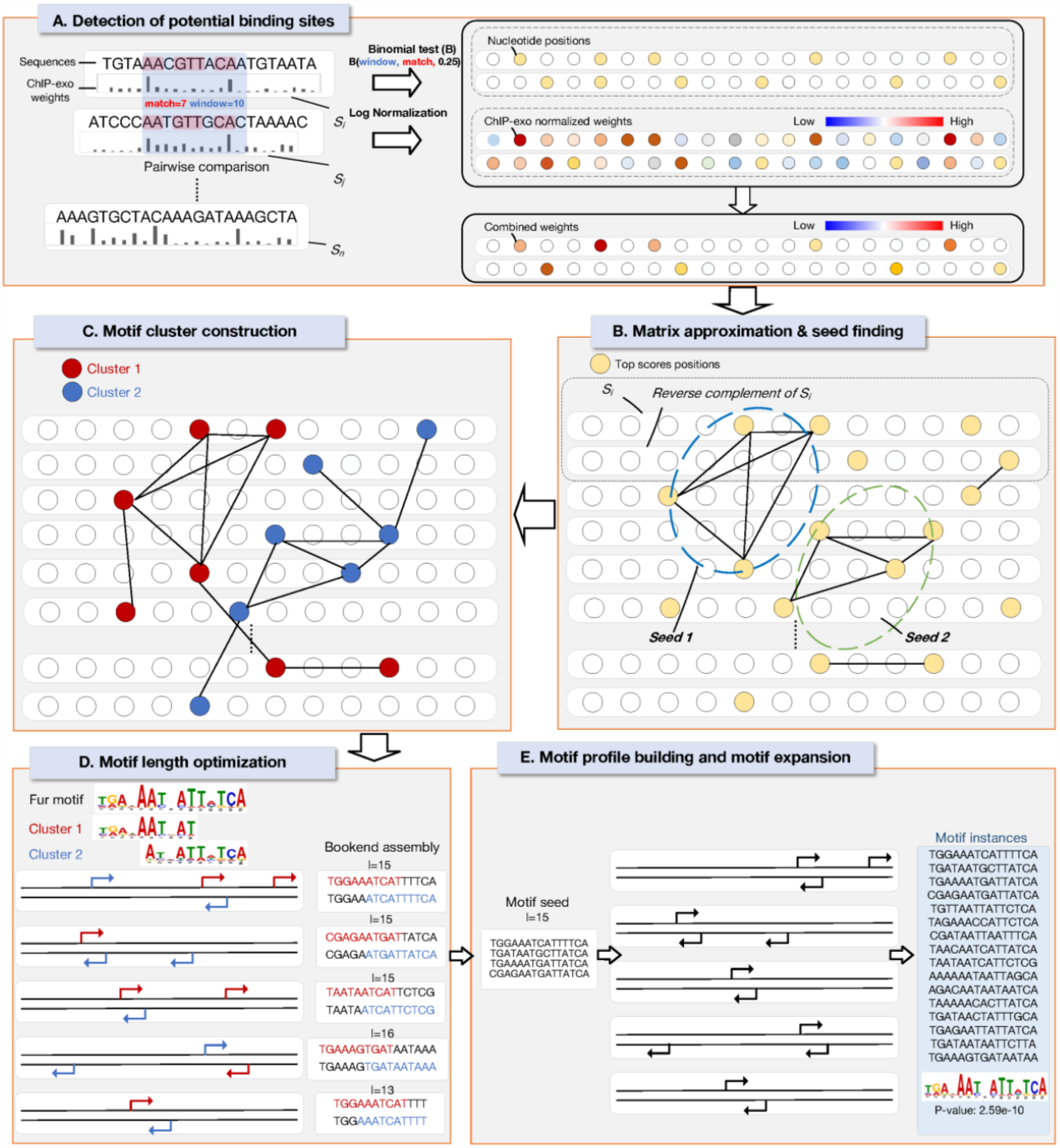
Schematic overview of TESA framework. TESA detects potential TFBSs from FASTA files by scoring each sequence segment based on binomial distribution **(A)**. Then, TESA constructs a graph by incorporating the highly ranked pairs of sequence segments **(B)**, and then performs clustering to identify motif seeds **(C)**. Afterwards, TESA optimizes lengths of motifs by relying upon a bookend model for motif seed assembly **(D)**. Finally, TESA refines the binding sites in each TF motif, yielding the ultimate motif profiles **(E)**.

### Data pre-processing

To obtain a nucleotide level weight score from ChIP-exo enriched regions using TESA, two input files are required, a reference genome file in FASTA format and a ChIP-exo read alignment file in SAM/BAM format^25^. TESA depends on two integrated toolkits, MACE and BEDTOOLS^23,24^, for peak calling and extraction of sequences as well as coverage score on each nucleotide of sequences, respectively. For the purpose of retaining more information, TESA selects flanking regions of 100 bp centered on each ChIP-exo peak by default, and assigns coverage score to each nucleotide. Finally, TESA normalizes the coverage score of each nucleotide, *x*_1_, *x*_2_, …, *x*_*n*_, as Eq. (1) to ensure that coverage scores on different peaks are on the same scale.

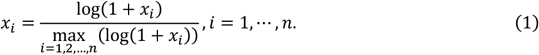

#### Step 1. Detection of potential binding sites

Without loss of generality, we suppose the input data has *m* sequences of length *n*, with each nucleotide assigned a normalized coverage score (**Figure 1A**). We then build a matrix *M*^*h*^, with size 2*m* × *n*. Rows of *M*^*h*^ represent the input peaks (odd rows) as well as their reverse complementary sequences (even rows); columns denote the starting positions of segments of length *l* on peaks; and values depict the normalized coverage scores. We identify potential TFBSs by two rounds of alignment of sequence segments, each of which demands an auxiliary matrix of the same size as *M*^*h*^, i.e., *M*^1^ and *M*^2^, respectively. Values in *M*^1^ and *M*^2^, which are initialized as zeros, denote the TF binding signals. TESA borrows the segment alignment steps from BoBro^18^. In the first round of alignment, for all segment pairs *s*_*ij*_ and *s*_*pq*_ of length *l* with *k* identical positions in the input peaks or their reversed complementary sequences, we calculate *f* and *f*^′^ as Eq. (2) and then store *f*^′^to *M*^1^,

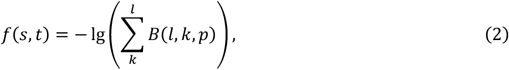

where *B*(·) is binomial distribution and *p* = 0.25, and

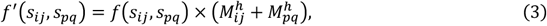

where 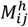 is the corresponding value of the start position of *s*_*ij*_ in matrix *M*^*h*^. We set *f* = 0 if 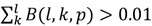. If either (*i*) *f*′ is among the top *t* (5 as default) of all pairs of sequence segment alignments between the two sequences, or (*ii*) *f* > 3, we add 0.5 (if the dinucleotides preceding *s*_*ij*_ and *s*_*pq*_ are identical) or 1 (else) to 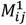 and 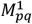, correspondingly. In the second round of alignment, we similarly calculate *f* and *f*^′^ based on *M*^1^, and then store the *f*^′^to *M*^2^ by

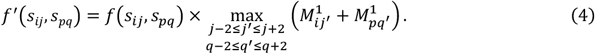

We normalize the matrices *M*^1^ and *M*^2^ using the same approach as that in data pre-processing. By sequence segment alignments, true signals as well as the flanking regions are enhanced (in both direction) in matrix *M*^2^, whereas false positive signals from ChIP-exo experiments and primary analysis will be suppressed. The three matrices act as the input to Step 2.

#### Step 2. Matrix approximation and seed finding

We update *M*^2^ by adding *M*^*h*^ on *M*^2^ and then build a graph, *G*, in which vertices represent all segments of length *l* in the input peaks and their reversed complementary sequences (**Figure 1B**). To connect vertices with edges, for each vertex pair in *G*, we calculate *f*′ by

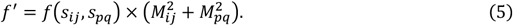

For each pair of segments, if *f*′ is among the top *t* (3 in default) among all pairs of segments between two sequences, connect *s*_*ij*_ and *s*_*pq*_ with an edge of weight *f*^′^. The graph *G* is input to Step 3 for clustering.

#### Step 3. Motif cluster construction

We identify clusters on *G* using a heuristic as follows (**Figure 1C**). First, we remove one-degree vertices, iteratively, until no one-degree vertex exists in *G*. Second, we construct a maximal independent set in G, i.e., no edge exists between any pair of vertices in it, by iteratively selecting the vertex with the largest degree. Third, we construct the seeds for graph clustering, which refer to the vertex set composed of one vertex in the maximal independent set with its neighbors. Fourth, we build clusters by linking the seeds that are connected in *G*. The set of clusters will be leveraged for motif length optimization in Step 4.

#### Step 4. Motif length optimization

To obtain the optimal lengths of motifs, we conduct a pairwise comparison among motif seeds and assemble the significantly overlapped pairs of clusters using a bookend model, detailed as follows (**Figure 1D**). For a pair of clusters, *c*_*i*_ and *c*_*j*_, composed of *n*_*i*_ and *n*_*j*_ vertices, respectively, we calculate the overlaps between each pair of vertices from *c*_*i*_ and *c*_*j*_ located on both input peaks and reverse complementary sequences with distance less than *d* (25 by default). Supposing *n*_*i*_ ≥ *n*_*j*_, we evaluate the statistical significance of the overlaps approximately by

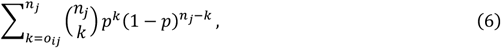

where *o*_*ij*_ is the number of matched nucleotides and *p* = (*dn*_*i*_)/(*mn*). The significantly overlapped pairs of clusters are kept, based on which the two clusters are concatenated by aligning the overlapped positions together and concatenating the sequence segments composing them. After this step, we obtain the collection of motifs to be input to Step 5.

#### Step 5. Motif profile building and motif expansion

After assembling significantly overlapped clusters, the length of the assembled motif is *l*_*p*_ = 2*l* − *l*_*o*_, where *l*_*o*_ is the number of overlapped nucleotides between sequence segments of the two clusters. Then, for each motif *p*, we build the preliminary motif profile, i.e., position weight matrix (PWM), as

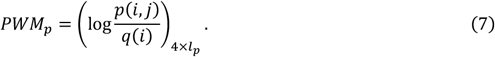

We calculate the similarity score, *S*(*s, p*), between *PWM*_*p*_ and a segment *s*_*i*_′_*j*_′ (with length *l*_*o*_) by

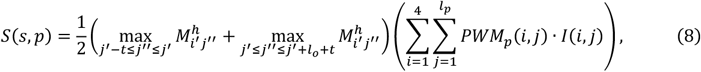

where *p*(*i, j*) is the probability of nucleotide type *i* appearing at position *j* in the alignment, and *q*(*i*) is the probability of *i* appearing in the simulated background sequences; *I*(*i, j*) indicates if the *j*-th nucleotide of *s*_*i*_′_*j*_′ is type *i*; *M*^*h*^ is the score which is defined in Step 1, and *t* is adjustable with default value 2. With this approach, we score the sequence segments not incorporated in the motif. If the similarity between *PWM*_*p*_ and a sequence segment is larger than that of at least one sequence segment included in the motif, we add the sequence segment to the motif. By following this process, we expand the set of motif instances and obtain a refined set of motifs (**Figure 1E**).

## RESULTS

### Benchmarking motif discovery on ChIP-exo datasets

We test the performance of TESA against three existing tools, which incorporate the ones considered to be the top-performing algorithms in the field of motif discovery from prokaryotes (BoBro, MEME-ChIP, and Weeder). The MEME-ChIP framework has adopted the STREME algorithm, which was recently published. The comparison of performance for sequence classification is evaluated by the Area Under Receiver Operating Characteristic curve (AUC), Accuracy (ACC), Matthews Correlation Coefficient (MCC), and area under the Precision-Recall Curve (PRC)^2,22^. We downloaded 20 ChIP-exo datasets of E. *coli* from the proChIPdb database^19^, which are composed of a wide array of peak numbers (25-1,630).

As showcased in the bar plots of **Figures 2A-2T**, TESA achieves the highest AUC values on 19 of the 20 datasets, with the exception of one dataset, on which MEME-ChIP surpasses TESA. BoBro showcases obvious advantages over MEME-ChIP and Weeder with higher AUCs than them on 12 datasets. This may be due to the fact that BoBro and TESA share a similar sequence segment alignment approach. On datasets composed of a relatively smaller number of sequences, TESA demonstrates particularly outstanding advantages. Similarly, MEME-ChIP is also apt to perform better on small datasets. This could be because MEME-ChIP has higher efficiency to optimize motif profiles on smaller datasets. It is also noteworthy that higher AUCs are observed for Weeder with the increase of dataset sizes. This is consistent with the fact that profiled-based methods, e.g., MEME, are more suitable for small datasets, while enumeration-based approaches perform better on relatively larger datasets. Regarding ACC, TESA, BoBro, and MEME-ChIP possess approximately the same performances across the 20 datasets. TESA concentrates upon enhancing and unraveling the signals (true positive) rather than filtering out the sequences without TFBSs, which may explain its average expression among the three algorithms, since ACC quantifies the capability to uncover both true positives and true negatives. We also calculate and compare the MCC score, which measures the difference between true TFBSs/non-TFBSs and false TFBSs/non-TFBSs for each of the four algorithms. TESA is ranked at the first place in 18 datasets, followed by BoBro and MEME-ChIP. Weeder does not perform well, as assessed by MCC. When it comes to PRC, TESA also occupies the top-ranked position on all datasets. Additionally, BoBro also perform well on nearly all datasets. This might be explained by the similar framework of TESA and BoBro.

**Figure 2.**
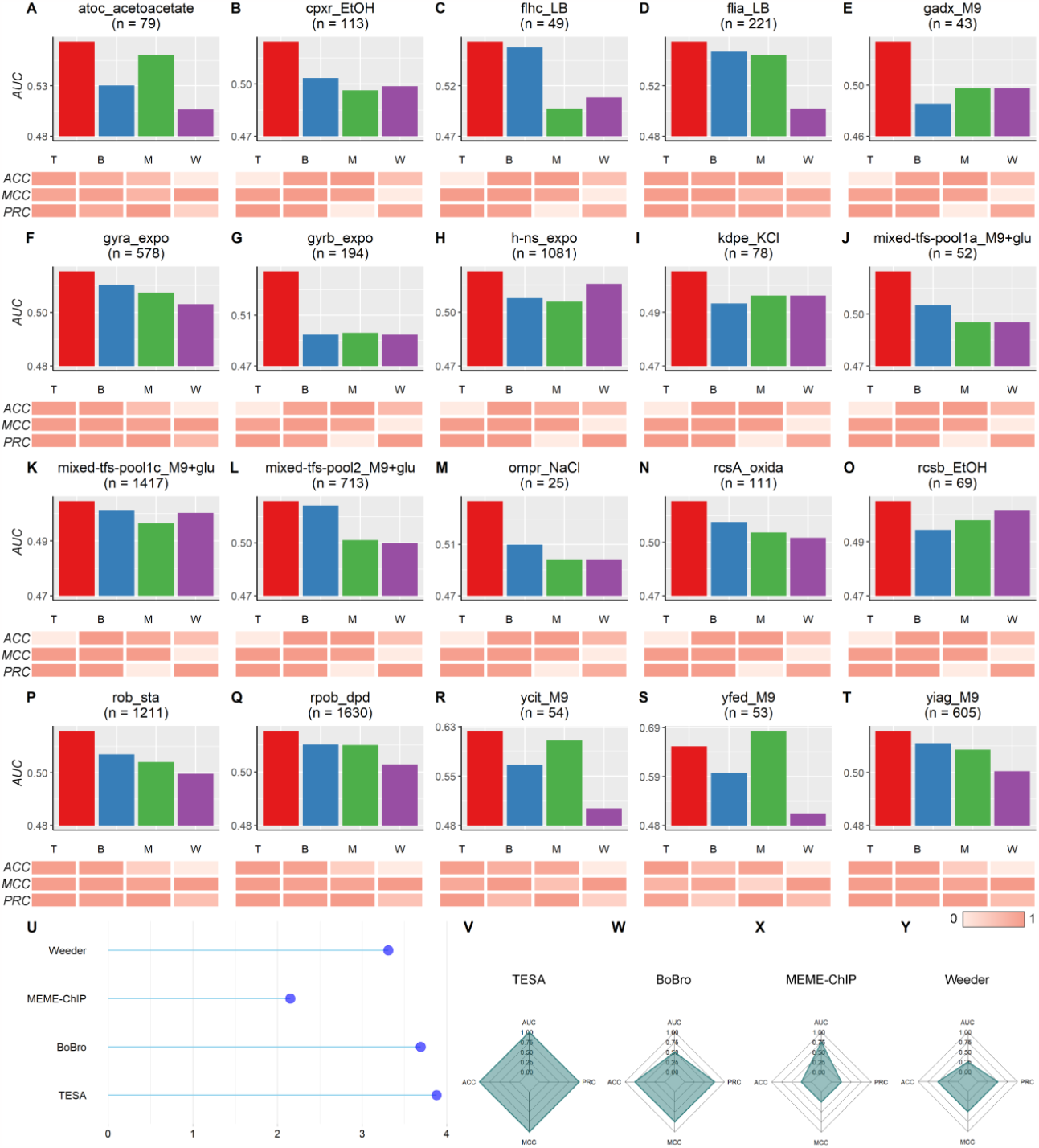
Overall performance of TESA. **(A-T)** Comparison of AUCs on each of the 20 datasets (bar plots) and relative performance of four algorithms regarding ACC, MCC, and PRC, respectively (heatmaps), which have been normalized over four algorithms T, B, M, and W represent TESA, BoBro, MEME-ChIP, and Weeder, respectively. **(U)** Overall scores of each algorithm summed across 20 datasets. **(V-Y)** Averaged scores of each algorithm in terms of AUC, ACC, MCC, and PRC.

To intuitively display the performance of an algorithm, we normalize and sum the four scores to obtain a unique one (**Figure 2U**). TESA and BoBro demonstrate better overall performance than Weeder and MEME-ChIP. In addition, we compare the performance of the four algorithms based on ranks of their scores. Specifically, a score of 4 is assigned to the algorithm that possesses the best performance, and a score of 1 is given to the method that has the worst performance. We calculated the average scores for each algorithm across the 20 datasets to gain insight into their overall performance and respective strengths and weaknesses (**Figures 2V-2Y**). TESA has the highest average score regarding each of the four scores, whereas BoBro also achieves high ACC, MCC, and PRC. Weeder and MEME-ChIP do not perform as well as TESA and BoBro. Similar to BoBro, Weeder also performs better, assessed by ACC, MCC, and PRC, compared with the evaluation using AUC. On the contrary, MEME-ChIP showcases relatively better performance evaluated using AUC, rather than ACC, MCC, and PRC. Generally, TESA is recommended among datasets, assessed by binary classification (AUC), ability to identify positive/negative sequences (ACC), correlation between predicted and actual labels (MCC), and capability of predicting positive sequences (PRC). BoBro and Weeder demonstrate complementary performance relative to MEME-ChIP.

### Comparison of predicted motifs with known motifs

To validate the discovered motifs, in this section, we compute the similarity between discovered motifs and the ones curated in DPInteract database by running TOMTOM and evaluate the statistical significance by *P*-values, *E*-values, and *Q*-values (**Table 1**)^20,21^. Due to the unavailability of motifs of some TFs in DPInteract, most of the motifs discovered by TESA show significant similarity (*Q*-values< 0.05) with that of TFs different from the ChIP-ed ones, which might be the co-factors of the ChIP-ed TFs regulating the same operons.

**Table 1.**
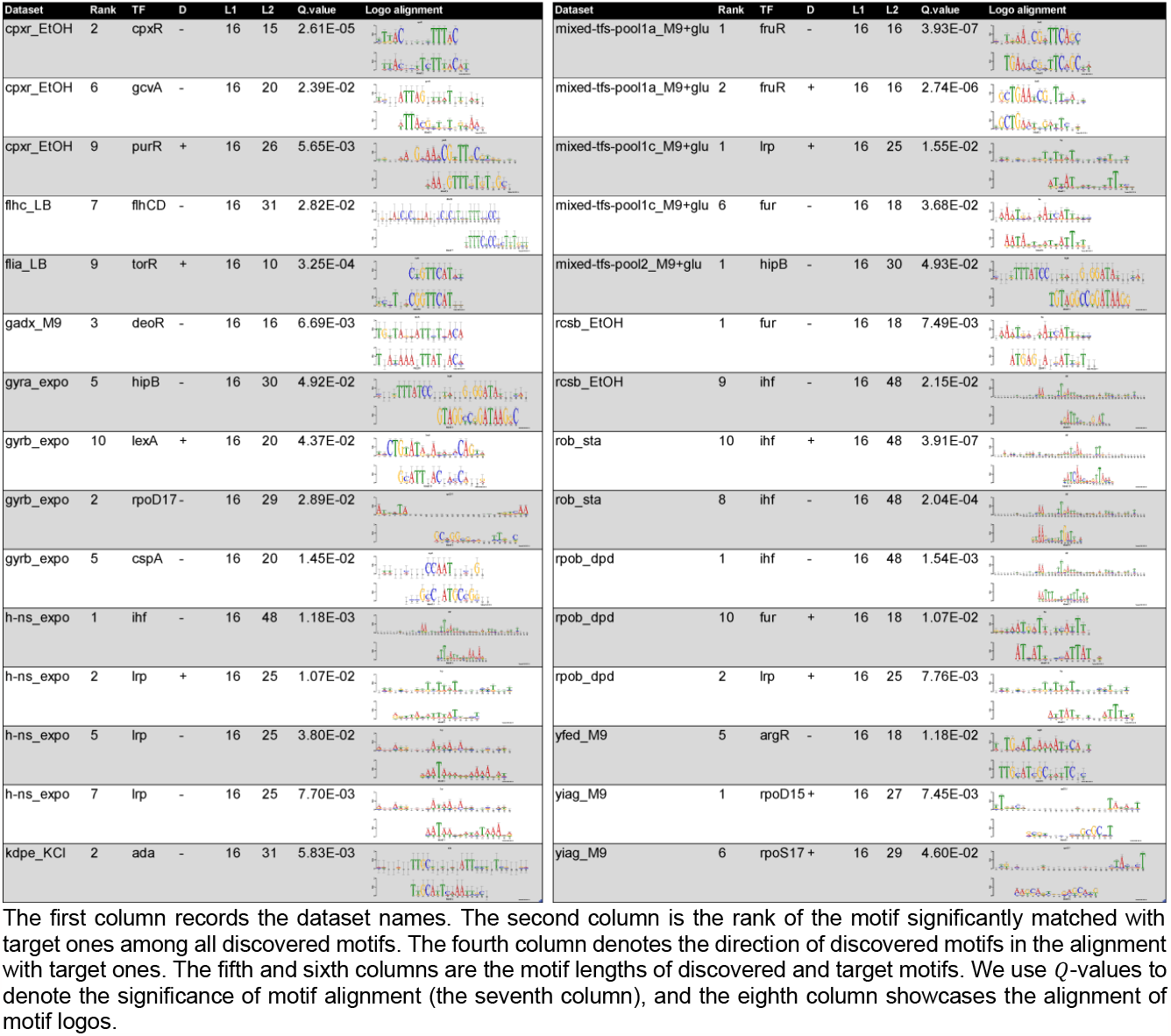
Similarity between target motifs and that discovered by TESA

## CONCLUSION

We develop a new algorithm TESA for prediction of TF motifs from ChIP-exo data of prokaryotic genomes, which improves the mainstream motif finding tools as we have shown in this article. Our performance analyses of TESA versus the other three programs suggest that the program can make accurate predictions of TFBSs evaluated by AUC, ACC, MCC, and PRC. Additionally, the motifs identified by TESA have high similarity to known ones in the DPInteract database. The reasons underlying the advantages of TESA may lie in: (*i*) incorporation of the ChIP-exo signals to score sequence segments, and (*ii*) efficient detection of motif lengths by merging significantly overlapped preliminary motifs. Although TESA possesses its strengths in motif discovery from ChIP-exo data, limitations exist, as follows: (*i*) alignment of sequence segments demand a large computational expense, and (*ii*) TESA depends upon multiple parameters in sequence segment alignment, which may not be applicable to data from other techniques. Therefore, our analysis results indicate a few directions for further improvement of the program. Specifically, we will consider (*i*) accelerating the sequence segment alignment step to improve efficiency; (*ii*) reducing the number of parameters which are defined in the algorithm, to make our program more generally applicable and usable for a broad array of ChIP-exo datasets.

## AVAILABILITY

The source code of TESA and a detailed tutorial can be found at https://github.com/Wang-Cankun/tesa.

## SUPPLEMENTARY DATA

Supplementary Data are available online.

## AUTHORS’ CONTRIBUTIONS

Q.M. and B.L. conceived the basic idea and designed the overall analyses. Y.L. implemented the experiments and completed the manuscript with the assistance of Y.W. C.W. developed the main program. A. F., A. M., J. J., and Z. L. proposed valuable suggestions on data analyses.

## ACKNOWLEDGEMENTS

The authors would like to thank Dr. Shuangquan Zhang for the assessment of motif finding programs.

